# Promotion of adipogenesis by JMJD6 requires the AT hook-like domain and is independent of its catalytic function

**DOI:** 10.1101/609982

**Authors:** Pablo Reyes-Gutierrez, Jake W. Carrasquillo-Rodríguez, Anthony N. Imbalzano

## Abstract

JMJD6 is a member of the Jumonji C domain containing enzymes that demethylate and/or hydroxylate substrate proteins. It is a multi-functional protein that has been implicated in disparate aspects of transcriptional and post-transcriptional control of gene expression, including but not limited to enhancer and promoter binding, release of paused RNA polymerase II, control of splicing, and interaction with the translation machinery. JMJD6 contributes to multiple aspects of animal development, including adipogenesis modeled in culture. We mutated proposed or characterized domains in the JMJD6 protein to better understand the requirement for JMJD6 in adipogenic differentiation. Mutation of JMJD6 amino acids that mediate binding of iron and 2-oxogluterate, which are required cofactors for enzymatic activity, had no impact on JMJD6 function, showing that catalytic activity is not required for JMJD6 contributions to adipogenic differentiation. In addition, we documented the formation of JMJD6 oligomers and showed that catalytic activity is not required for oligomerization, as has been reported previously. We also observed no effect of mutations in the sumoylation site and in the poly-serine stretch. In contrast, mutation of the AT hook-like structure, which mediates interaction with DNA and/or RNA, compromised JMJD6 function. The ability of JMJD6 to interact with nucleic acids may be a critical requirement for its function in adipogenic differentiation. The requirement for the AT hook-like domain and the lack of requirement for catalytic activity giving rise to the idea that JMJD6 may be functioning as a scaffold protein that supports the interactions of other critical regulators.

## Introduction

The Jumonji (Jmj) family of proteins encodes evolutionarily conserved oxygenases dependent on ferrous iron and 2-oxogluterate to hydroxylate metabolites, proteins and nucleic acids (1, 2). The conserved JmjC domain is structurally related to the cupin domains found in archaea and other kingdoms that possess active sites containing a metal ion within a histidine cluster (3). JmjC domains form a double-stranded β-helical fold in which eight β-strands form two, four-stranded antiparallel β-sheets (4). Distinctions between different JmjC families are generally defined by the structural elements that surround the conserved JmjC domain and the presence of other protein domains, many of which are interaction surfaces for chromatin or chromatin-bound proteins. The JmjC proteins themselves are overwhelmingly described as factors that promote the regulation of transcription and/or chromatin (5).

Phylogenetically, the JMJD6 protein belongs to the JmjC subfamily of hydroxylase enzymes (5). It has functions in myriad processes, including regulation of transcription, post-transcriptional control, splicing, local chromatin structure, and genome integrity (6–8). It has also been reported to be a secreted protein that is part of the extracellular matrix (9). Mechanisms of action vary widely. JMJD6 can bind chromatin, and it regulates transcription via enhancer and promoter binding as well as via regulation of elongation (10–14). JMJD6 also binds RNA and multiple proteins involved in splicing, RNP formation, and mRNA export (15–21). The list of proteins that can interact with JMJD6 is large and continues to grow (Kwok review). Not surprisingly, JMJD6 has been proposed as a driver of multiple types of cancer through several of its different functions (13, 22–33).

Knockout of *Jmjd6* in mice resulted in normal development until E12.5, but pleiotropic developmental phenotypes were observed by E15.5. These included craniofacial and cardiac malformations, blocked or delayed lung, intestine, erythropoietic, and immune cell differentiation, as well as subcutaneous edema. *Jmjd6* deficient mice died pre- or peri-natally (34–37). The role of JMJD6 in development is poorly understood, though there are multiple lines of evidence that it may play a role in apoptosis regulation (7). Morpholino-induced knockdown in zebrafish resulted in altered embryonic cell migration, with the frequency and extent of developmental deficiencies and death showing a linear response to the amount of morpholino used (38). Recently, JMJD6 was shown to mediate body axis patterning in *Xenopus* through transcriptional regulation of the Tcf7l1 repressor protein (14). We previously demonstrated a requirement for JMJD6 in promoting differentiation of adipocytes by two distinct mechanisms: (1) promoter binding and transcriptional activation of the PPARγ and C/EBPα master regulators of adipogenesis and (2) a post-transcriptional mechanism that elevated the levels of the C/EBPβ and δ proteins immediately after the onset differentiation signaling (11).

Presumably, JMJD6 function is tied to its enzymatic activity, but the nature of this activity remains controversial. JMJD6 was first reported to be a histone arginine demethylase (39), but this result has been questioned (17, 19, 40). Nevertheless, subsequent studies expanded the range of substrates for JMJD6 demethylation (12, 18, 27, 41–44). Other work indicates that JMJD6 is an RNA demethylase (12) as well as a lysyl oxidase that modifies a range of substrates (19, 20, 24, 40, 45, 46). Finally, a recent report describes JMJD6 as a kinase capable of phosphorylating histone H2A.X (33). Despite the myriad possibilities for enzymatic activity, JMJD6 can also act in a manner independent of its known enzyme functions. JMJD6 cooperates with U2AF65, a necessary accessory factor in the splicing process (47, 48) co-regulate alternative splicing (19, 21). JMJD6 lysyl hydroxylase activity was required for some, but not all, alternative splicing events in *Jmjd6* null embryonic tissues (21). These findings provide some resolution to the conclusions of prior studies where JMJD6 enzymatic function was reported to be required (19) or not (16) for alternative splicing. In other work, knockdown of JMJD6 inhibited adipocyte differentiation, which could be rescued by exogenous expression of either wild type JMJD6 or a catalytically inactive JMJD6 protein (11). Thus the contributions of JMJD6 to adipocyte differentiation may be independent of enzymatic activity.

In this study we further explored the requirement for JMJD6 catalytic activity in promoting adipogenesis and evaluated the role of other domains of the JMJD6 protein. Mutation of JMJD6 amino acids that permit interaction with iron, with the required cofactor, 2-oxogluterate, or with the substrate had no effect on the ability of JMJD6 to rescue defective adipogenesis caused by JMJD6 knockdown. Similar results were obtained for mutation of the sumoylation site or mutation of the serine-rich domain in the C terminal portion of the protein. In contrast, mutation of the AT hook-like domain motif did compromise rescue. This work identifies the AT hook-like domain as a critical domain through which JMJD6 promotes adipogenesis.

## Results

We previously examined the role of JMJD6 and determined it promoted adipogenesis in culture through both transcriptional and post-transcriptional mechanisms (11). In that study, we showed that shRNA-mediated knockdown of *Jmjd6* inhibited adipogenesis and that ectopic expression of JMJD6 could rescue the differentiation deficiency. We also rescued the shRNA-induced deficiency in differentiation by expressing a JMJD6 protein mutated at amino acids 187 and 189, which are critical for iron binding (11). Others have shown that mutations in one or both of these amino acids significantly compromise or abolish JMJD6 enzymatic activity (10, 24, 39, 40, 43, 45). Our results suggest that the catalytic activity of JMJD6 is not necessary for its role in adipogenesis and raise the question of which JMJD6 domains are required for this function.

The JMJD6 protein contains multiple predicted and characterized domains (Fig. 1A). The JmjC domain is responsible for enzymatic activity and contains amino acids that are important for binding iron and 2-oxogluterate, essential cofactors for enzymatic activity (discussed above and reviewed in (5)). A region that interacts with BRD4, a transcriptional regulator affecting RNA polymerase II elongation (reviewed in (49–51)), exists in the N-terminal portion of the JMJD6 protein (52). An AT hook-like domain is present C-terminal to the JmjC domain. Classical AT hooks are DNA interacting structures promoting interaction with the minor groove of DNA (53). A related structure, called the extended AT hook domain is somewhat different. An extended AT hook domain is extended in both the C- and N-terminal directions by basic amino acids, which gives the domain higher affinity for RNA than for DNA (54). JMJD6 contains a recognizable classic AT hook with additional basic amino acids in the C-terminal, but not the N-terminal, direction. We will refer to the JMJD6 domain as an AT hook-like domain. A poly-serine stretch exists in the C-terminal portion of JMJD6. This domain may be responsible for regulating the partitioning of JMJD6 between the nucleoplasm and nucleolar compartments (55). JMJD6 is predicted to be a target for sumoylation on K317 (56). Sumoylation is the post-translational addition of small ubiquitin-like modifier proteins to a target protein that can affect the regulation of gene expression, regulation of cell cycle, protein stability, protein transport (reviewed in (57–59)). A nuclear export signal is also present in the C-terminal part of the molecule, and at least five nuclear localization signals have been identified, three of which have been experimentally validated (56, 60).

**Figure 1.**
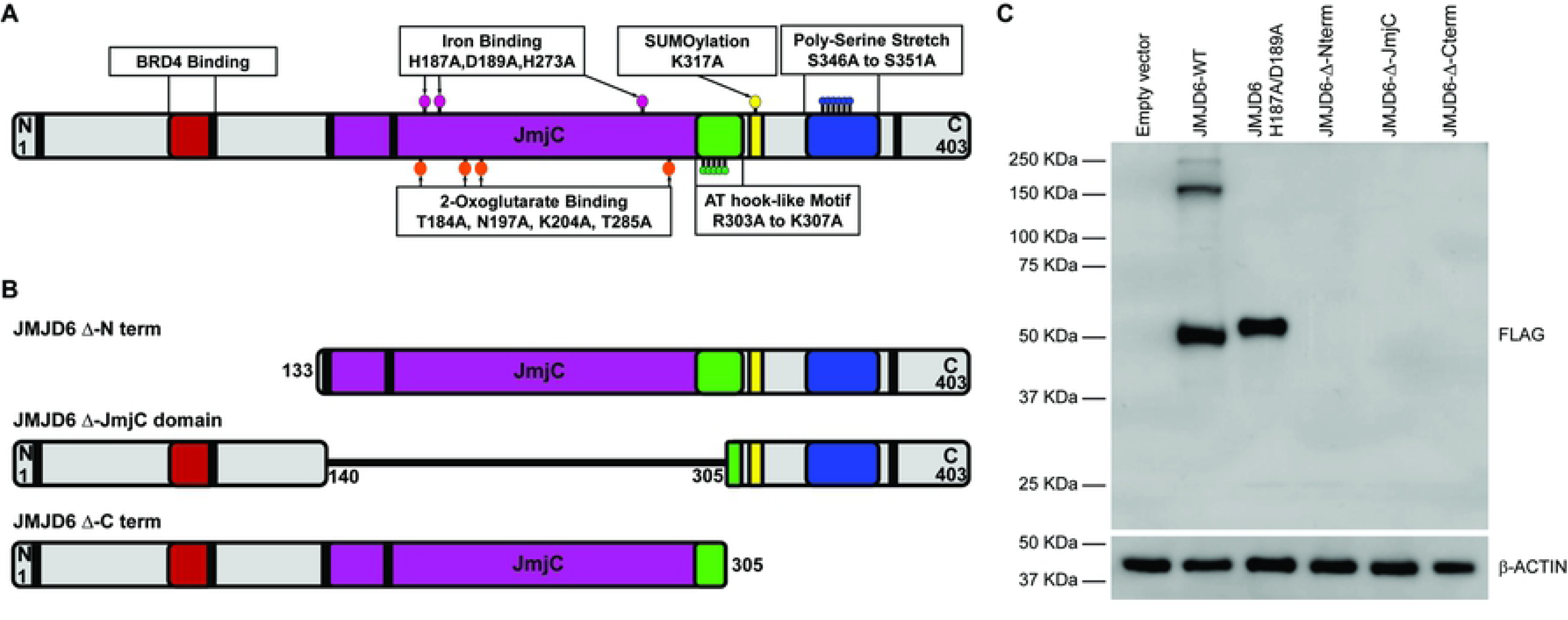
JMJD6 domains. **(A)** Schematic representation of JMJD6 domains. Mutations of specific amino acids are shown as colored ballons. Amino acids related to Fe^2+^ binding (His187, Asp189, His273), 2-oxogluterate binding (Thr184, Asn197, Lys204,Thr285), AT hook like motif (Arg303 to Lys307) Sumoylation site (Lys317) and Poly-serine stretch (Ser346 to Ser351) were substituted with alanine. Black rectangles represent nuclear localization sequences. **(B)** Representation of truncated mutants generated by deletion of the amino, JmjC or carboxy terminal regions of JMJD6. **(C)** Representative western blot of FLAG-tagged JMJD6 ectopically expressed in C3H10T1/2 cells. Cells selected for the expression of the resistance gene encoded by the retrovirus expressed either wild type or catalytically inactive JMJD6 H187A/D189A but did not express the truncated mutants. The expected sizes of truncated JMJD6 were 32 kDa (Δ Nterm), 28 kDa (Δ JmjC), and 35 kDa (Δ Cterm). β-ACTIN expression was monitored as a control.

We and others have had technical issues expressing JMJD6 deletion mutants in eukaryotic cells. Although two groups were able to express JMJD6 deletion mutants in HEK293 cells (12, 40), others were only able to achieve expression of some deletion mutants in HEK293 cells (61). We previously reported that we were unable to express a series of deletion mutants encoded within lentiviral vectors in mouse multipotent mesenchymal C3H10T1/2 cells (11). The reasons for this difficulty are unclear. We subsequently attempted to express three different deletion mutants – one lacking sequences N-terminal to the JmjC domain, one lacking sequences C-terminal to the JmjC domain, and one lacking the JmjC domain (Fig. 1B). Cells showing expression could not be obtained (Fig. 1C). Consequently, we resolved to limit the mutagenesis strategy to create point mutations in one or in a small number of amino acids.

### Mutation of amino acids critical for cofactor binding and catalysis has no impact on adipogenesis

Previously we showed that different shRNAs targeting *Jmjd6* in C3H10T1/2 cells prevented differentiation into lipid containing adipocyte-like cells (11). Here, we reiterate those findings by showing that cells expressing shRNA targeting the 3’ untranslated region of *Jmjd6* and an empty expression vector had reduced JMJD6 protein levels whereas cells expressing shRNA targeting *Jmjd6* and an expression vector expressing a FLAG-tagged *Jmjd6* cDNA restored JMJD6 levels to those observed in cells expressing a scrambled sequence control shRNA (Fig. 2A; compare lane 1 showing scrambled shRNA control to lanes 2 and 3). Note that JMJD6 forms stable oligomers despite electrophoresis in the presence of reducing agent in a denaturing gel, as has previously been observed (55, 61, 62). The significance of JMJD6 oligomerization is not well understood. As expected, cells with reduced levels of JMJD6 expressing empty vector showed an inhibition of differentiation, while expression of the FLAG-tagged JMJD6 restored differentiation, as demonstrated by Oil Red O staining of accumulated lipids in the cells (Fig. 2B; compare panels 1-3). When the JMJD6 protein containing mutations in the iron binding amino acids H187A/D189A was expressed, or when another amino acid mediating iron binding (H273; (17, 45)) was mutated and expressed, the differentiation deficiency caused by the shRNA targeting *Jmjd6* was rescued (Fig. 2B, panels 4-5). Consistent with prior results (40), expression of the mutations in the iron binding sites restored levels of the 50 kD monomer form of JMJD6 but prevented oligomerization for reasons that are not known (Fig. 2A, lanes 4-5).

**Figure 2.**
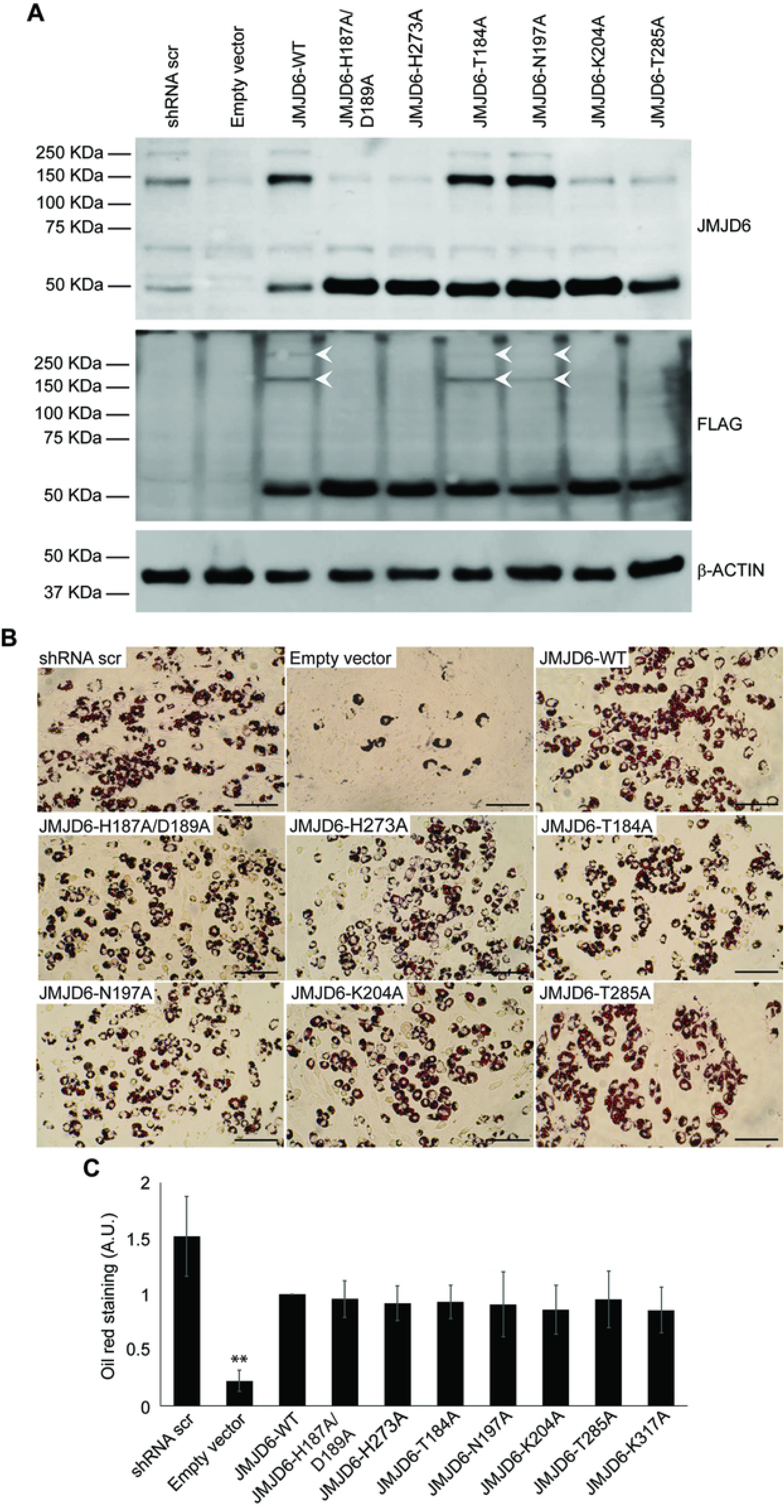
Expression of catalytically deficient JMJD6 mutants rescues the adipogenic differentiation deficiency caused by JMJD6 knockdown. **(A)** Representative western blots for JMJD6 or ectopically expressed JMJD6 (FLAG) in C3H10T1/2 cells with stable knockdown of JMJD6. Controls included cells expressing a scramble shRNA that does not affect JMJD6 expression and JMJD6 knockdown cells expressing the empty vector (lanes 1-2). White arrows indicate multimerization of expressed mutants. β-ACTIN levels were monitored as a control. **(B)** Representative Oil Red O staining of C3H10T1/2 cells with stable expression of either scramble shRNA or shRNA against *Jmjd6* that were expressing the empty vector or the wild type or the indicated JMJD6 mutants. Staining was performed after 6 days of differentiation. Scale bar = 100µm. **(C)** Quantitation of Oil Red O staining. The values are the average of optical density at 500nm of three independent experiments. The data are relative to the value for the JMJD6 wild type sample, which was normalized to 1. **P<0.01; A.U. – arbitrary units.

We then created point mutations (T184A, N197A, K204A, T285A) that, based on the crystal structure of JMJD6 and other 2-oxogluterate-dependent oxygenases (45, 56, 63–66) are implicated in the binding of 2-oxogluterate, an obligate cofactor for catalysis. T285 may also play a role in substrate recognition (45). K204A and T285A have been shown to be inactive for hydroxylation (45). Each of the four mutants restored expression of the 50 kD JMJD6 monomer, but only two (T184A, N197A) restored expression of the oligomeric forms of JMJD6 (Fig. 2A, lanes 6-9). Despite these differences, all four mutants were able to support adipogenic differentiation (Fig. 2B, panels 6-9). Oil Red O quantification from multiple experiments is shown in Fig. 2C. The data clearly demonstrate that binding of cofactors required for JmjC domain catalysis is not required to promote adipogenic differentiation, and that substrate recognition is also likely not required. In addition, these data demonstrate that there is no correlation between the ability of JMJD6 to bind to critical enzyme cofactors required for catalysis and the ability of JMJD6 to multimerize.

### The sumoylation site on JMJD6 is not required for adipogenic differentiation

JMJD6 is predicted to be sumoylated on the lysine at amino acid position 317 (56). Mutation of K317 to alanine restored expression of the JMJD6 monomer, did not affect the formation of JMJD6 oligomers, and restored adipogenic differentiation to cells expressing shRNA targeting *Jmjd6* (Fig. 3A-B). Quantification of Oil Red O staining in the presence of the K317A mutation is shown in Fig. 2C. We conclude that sumoylation of JMJD6 is not required for adipogenic differentiation.

**Figure 3.**
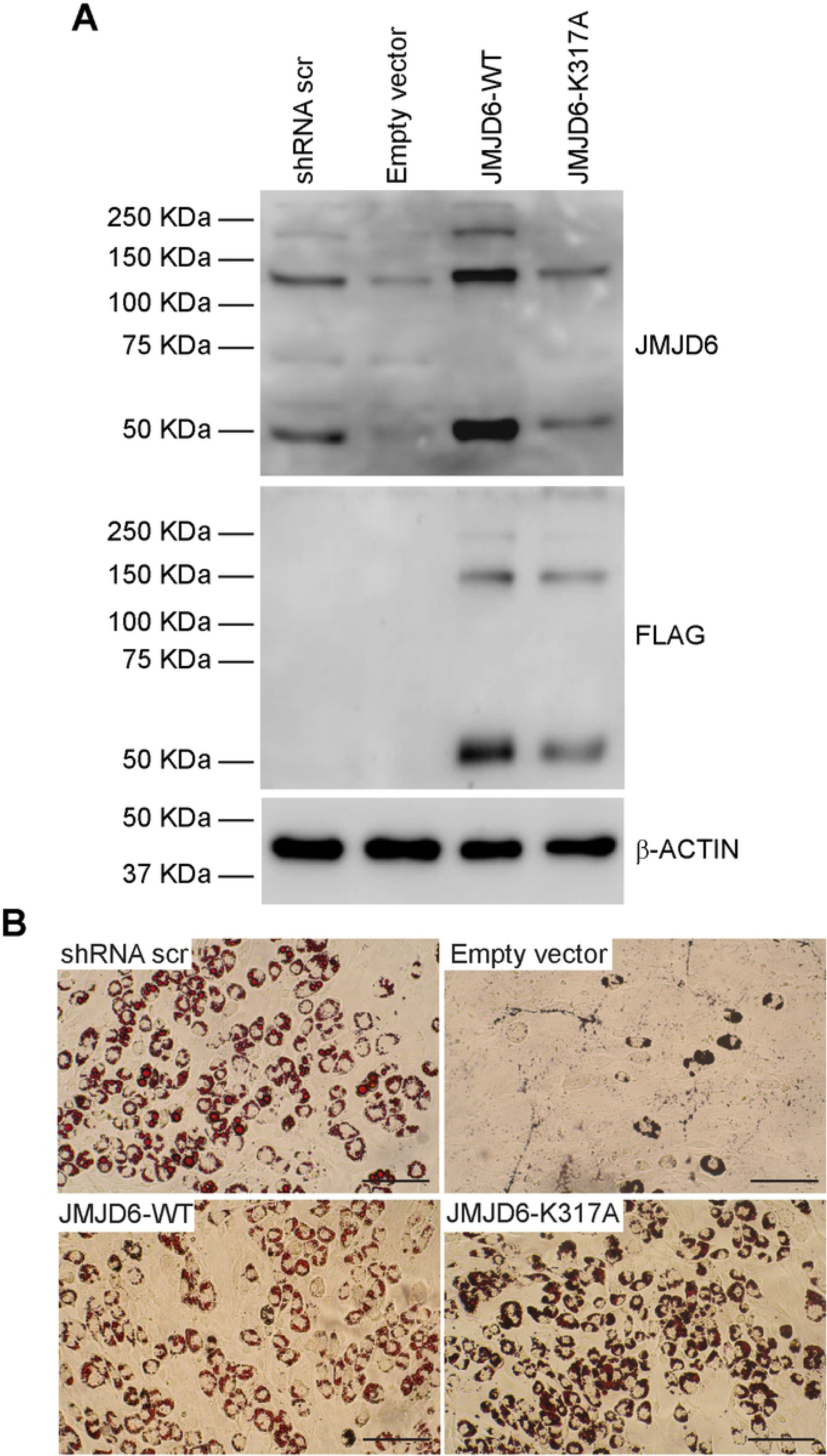
The predicted sumoylation site in JMJD6 is not essential for adipogenesis. **(A)** Representative western blots for JMJD6 or ectopically expressed JMJD6 (FLAG) in differentiated cells with stable knockdown of JMJD6. Controls included cells expressing a scramble shRNA that does not affect JMJD6 expression and JMJD6 knockdown cells expressing the empty vector (lanes 1-2). β-ACTIN levels were monitored as a control. **(B)** Representative Oil Red O staining of differentiated cells. Scale bar = 100 µm. Quantification of Oil Red O staining is presented in Figure 2C.

### Cells expressing JMJD6 point mutations in the BRD4 binding domain could not be propagated

JMJD6 has a role in controlling elongation by RNA polymerase II in conjunction with the BRD4 transcriptional regulatory protein. BRD4 mediates recruitment of JMJD6 to distal enhancers that counteract pausing by RNA polymerase II via long-range interactions (12). Targeting the bromodomain and extra-terminal domain (BET) protein family, which includes BRD4, with an inhibitory drug interferes with the ability of JMJD6 to transcriptionally activate the PPARγ and C/EBPα master regulators of adipogenic differentiation (11). BRD4 has also been shown to bind with lineage-determining transcription factors at enhancer sequences that are active during adipogenesis and myogenesis to promote lineage-specific gene expression (67, 68). Structural studies of the JMJD6-BRD4 association identified an α-helix formed by JMJD6 amino acids 84-96 that interacts with the extra-terminal domain of BRD4 (52).

We created three mutant alleles in the BRD4 interaction domain. One contained three substitutions (L90A, K91A, R95A), while another substituted alanine for each of the JMJD6 amino acids from 90 to 96. Expression of these mutant JMJD6 proteins was observed after the initial selection of lentiviral infected cells (Fig. 4), but expression was invariably extinguished within two-three subsequent passages despite repeated generation of expressing cells. Consequently, the contributions of the BRD4 interacting domain toward adipogenesis could not be assessed. A third allele contained alanine substitutions for each of the JMJD6 amino acids from 84 to 89, but was never detected as an expressed protein in any experiment.

**Figure 4.**
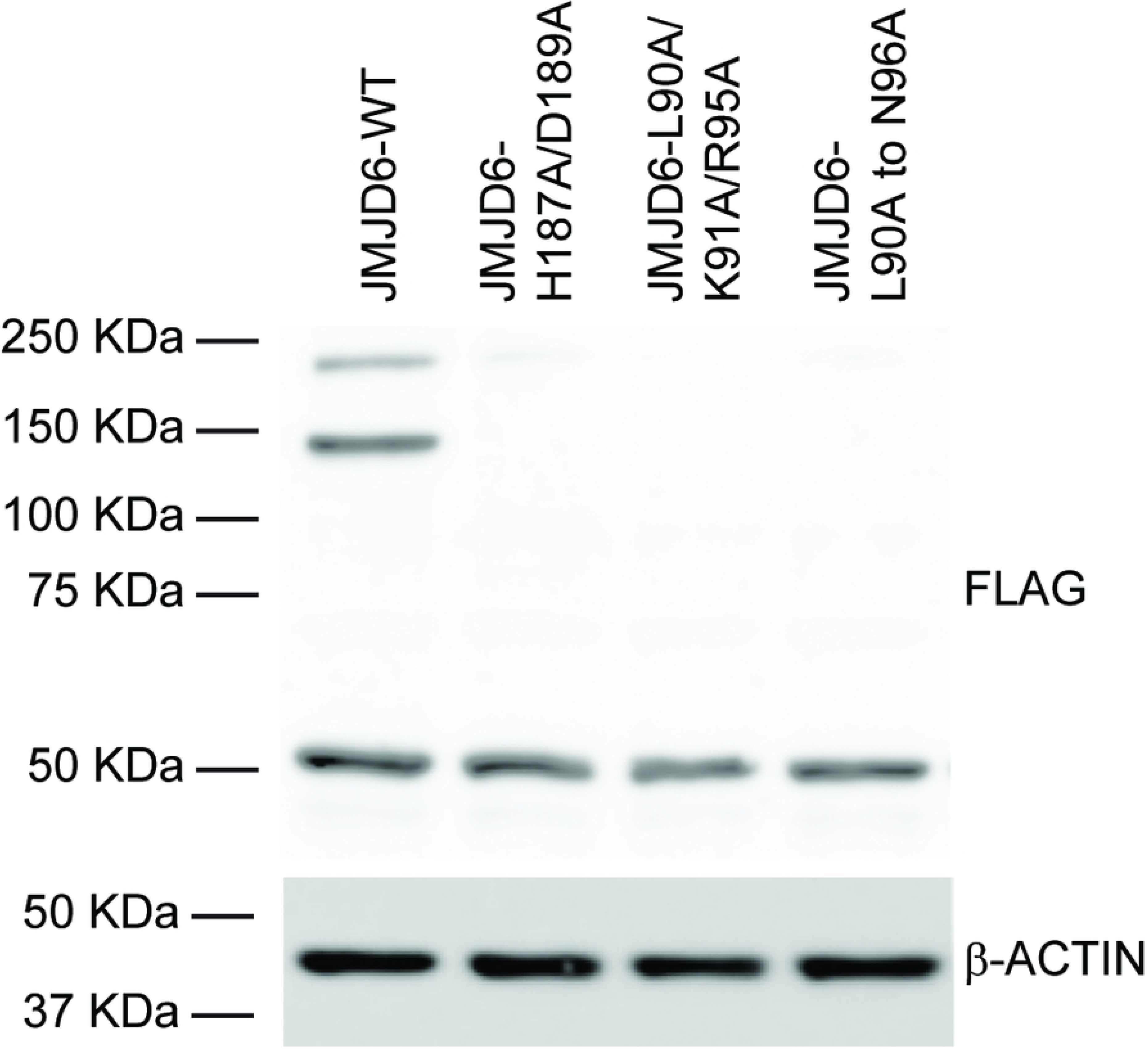
Expression of JMJD6 mutated in the BRD4 binding domain. Representative western blot showing that expression of FLAG-tagged mutants JMJD6-L90A/K91A/R95A and JMJD6-L90A to N96A was detected only immediately after transduction and selection. Expression of the JMJD6 wildtype and the iron binding mutant H187A/D189A are shown as controls. β-ACTIN levels were monitored as a loading control.

### The JMJD6 AT hook-like domain, but not the poly-serine stretch, is required for adipogenesis

We replaced JMJD6 amino acids 303-307 or JMJD6 amino acids 346-351 with alanines to target the AT hook-like domain or the poly-serine stretch, respectively. Western blot analysis showed that both proteins could be expressed but the poly-serine stretch mutant was unable to form oligomers (Fig. 5A). Oil Red O staining of differentiated cells expressing the AT hook-like domain mutant was reduced, while there was no effect of the mutation in the poly-serine stretch (Fig. 5B). The result was confirmed by quantification of Oil Red O staining (Fig. 5C). Since one mechanism of JMJD6 action is to promote the expression of the adipocyte lineage determining transcription factors (Hu 2015), we examined PPARγ2 and C/EBPα mRNA expression in control cells and cells expressing the mutations in the AT hook-like domain or the poly-serine stretch. Expression of PPARγ2 and C/EBPα was compromised in the cells expressing the AT hook-like domain but not in cells expressing the mutation in the poly-serine stretch (Fig. 5D). This result provides a molecular explanation for the failure of the AT hook-like mutant to differentiate. We conclude that the AT hook-like domain contributes to JMJD6 function in promoting adipogenesis.

**Figure 5.**
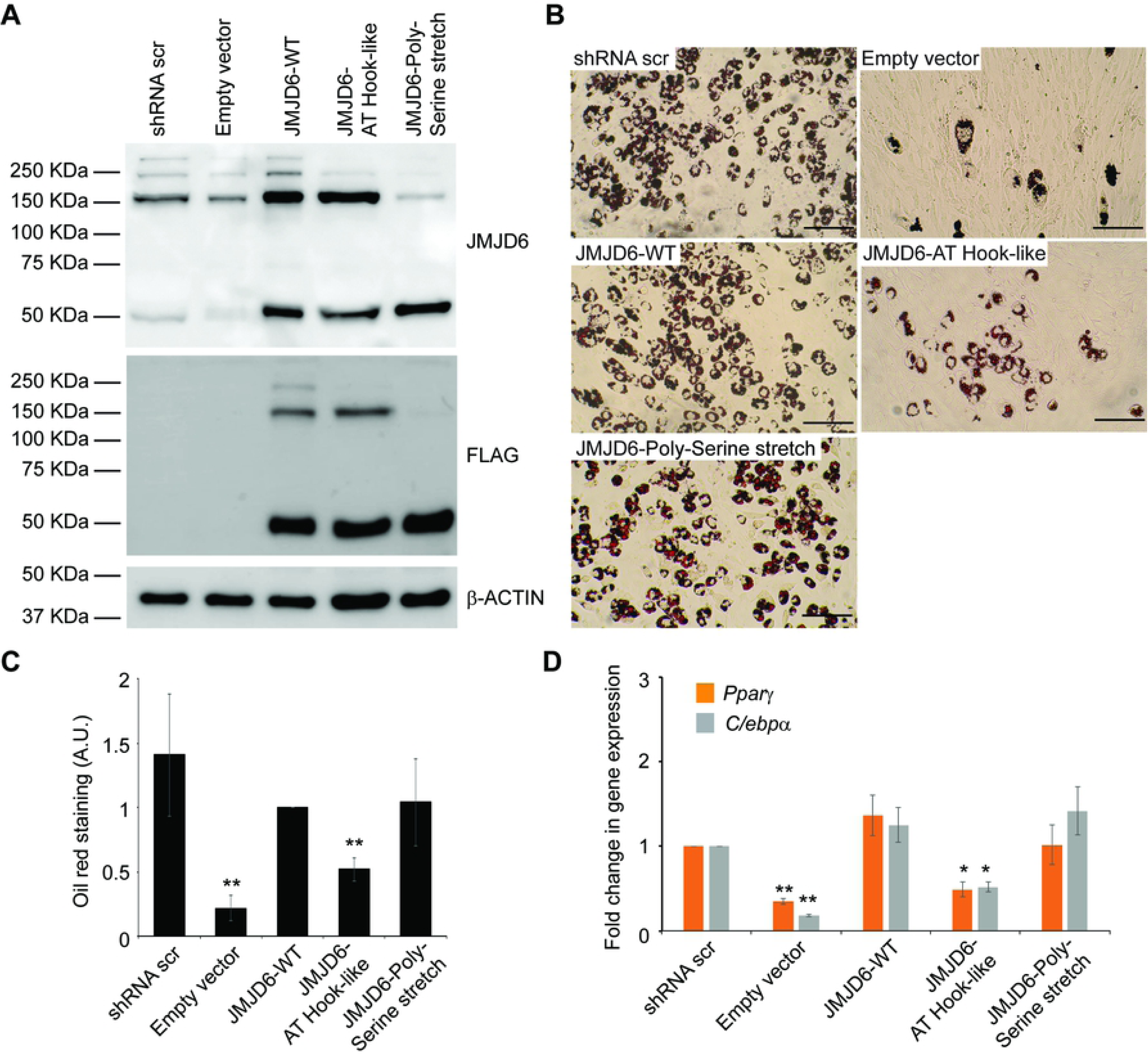
The AT hook-like domain of JMJD6 contributes to adipogenesis. **(A)** Representative western blots for JMJD6 or ectopically expressed JMJD6 (FLAG) in differentiated cells with stable knockdown of JMJD6. Controls included cells expressing a scramble shRNA that does not affect JMJD6 expression and JMJD6 knockdown cells expressing the empty vector (lanes 1-2). β-ACTIN levels were monitored as a control. **(B)** Representative Oil Red O staining of differentiated cells with stable expression of either scramble shRNA or shRNA against *Jmjd6* that were expressing either the empty vector, or the wild type, or the indicated JMJD6 mutants. Staining was performed after 6 days of differentiation. Scale bar = 100 µm. **(C)** Quantitation of Oil Red O staining. The values are the average of optical density at 500nm of three independent experiments. The data are relative to the value for the JMJD6 wild type sample, which was normalized to 1. **P<0.01; A.U. – arbitrary units. **(D)** Analysis of mRNA levels of the adipocyte lineage-determinant genes, *Cebpα* and *Pparγ*, in differentiated JMJD6 knockdown cells expressing wild type JMJD6 or the JMJD6-Polyserine stretch or JMJD6-AT hook-like mutants. The individual mRNA levels were normalized to *Eef1a1* mRNA levels. The normalized expression levels of shRNA scramble control cells were set as 1. *P<0.05 **P<0.01.

We used PyMOL to visualize the JMJD6 structure (PDB ID 3K2O; Mantri, 2010, JMB) and model the substitutions we made in the JMJD6 AT hook-like domain. Fig. 6 shows the JMJD6 dimer structure with the AT hook-like domain indicated and an isolated view of the expected structure (right) and the structure predicted when amino acids 303-307 (GPRK) are replaced by alanines (left). The substitutions affect the hinge between two α-helices but do not appear to significantly alter the positions of the α-helices. Instead the major change appears to be the absence of the side chains of the RPK amino acids, which may, directly or indirectly, affect the predicted interaction with RNA (17, 54).

**Figure 6.**
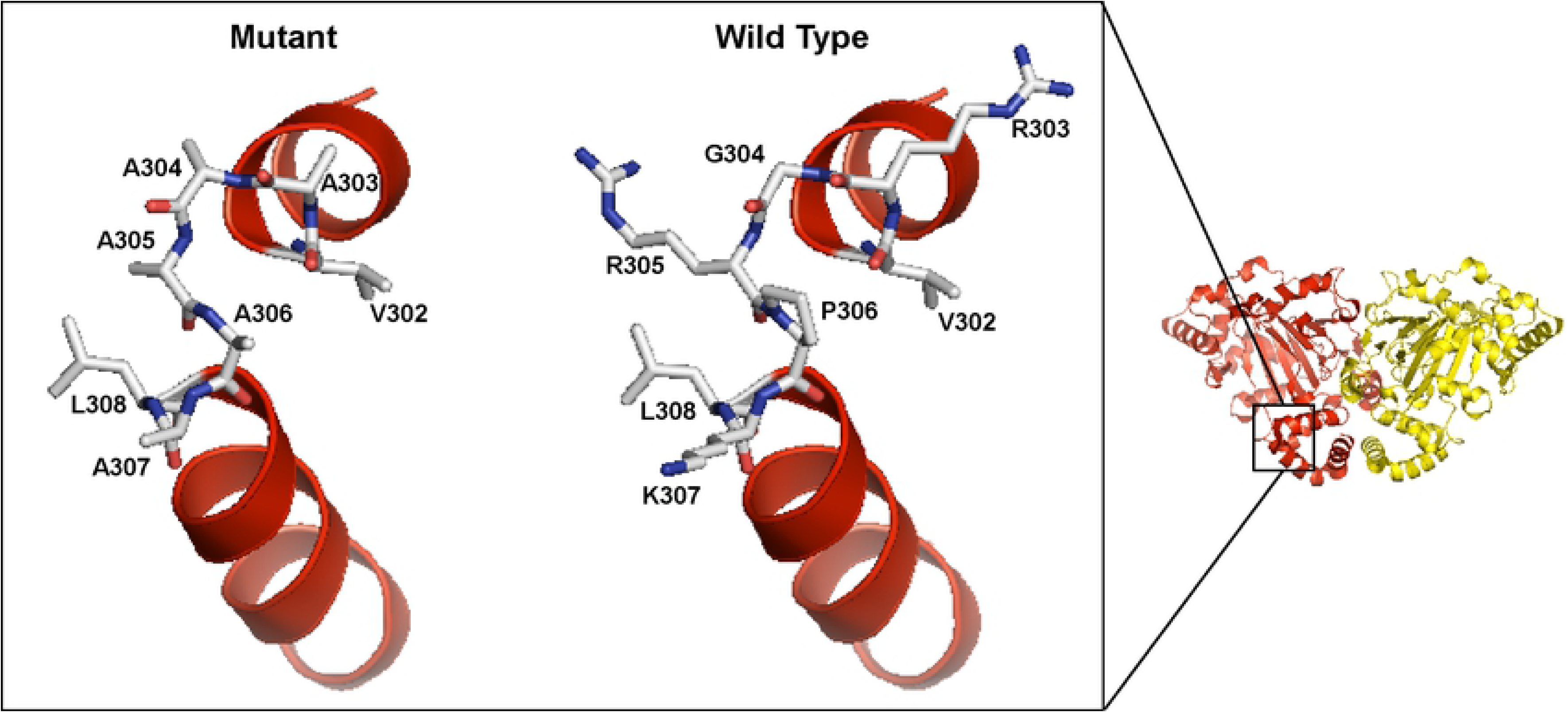
Predicted structure of the JMJD6 AT hook-like domain. PyMOL was used to visualize the JMJD6 dimer structure reported by Mantri et al (PDB ID 3K2O; (45)). The magnified area is a comparison between the predicted hinge area of the wildtype motif (right) and the motif with alanine substitutions of the GPRK amino acids at positions 303-307 (left).

## Discussion

### JMJD6 enzymatic activity is not required for promoting adipogenesis in culture

Three JMJD6 amino acids (H187, D189, H273) combine to form an iron binding site (17, 45). Multiple reports have demonstrated that mutation of these amino acids, alone or in combination, blocks arginine demethylase and/or lysyl hydroxylase enzymatic activities (19, 39, 45). We previously published that expression of the double mutant H187A/D189A rescued the differentiation deficiency in cells expressing shRNA targeting *Jmjd6* to the same extent as did expression of a wild type JMJD6 protein (11). The implication of the prior finding is that all aspects of JMJD6 function in the adipogenic differentiation process are independent of iron binding, and hence, catalytic activity. Here we further support the prior finding by showing that expression of the H273A mutation also has no impact on adipogenic differentiation.

JmjC family proteins also bind to 2-oxogluterate, which catalyzes oxidation reactions in coordination with iron. 2-oxogluterate is an obligatory cofactor for catalysis. The crystal structure of JMJD6 predicts hydrogen bonding of T184, N197, and K204 with 2-oxogluterate in the 2-oxogluterate binding pocket (45). Thr285 is important for the positioning of N197 (45). To further investigate the requirement for JMJD6 catalysis in adipogenesis, we created single point mutations in JMJD6 amino acids that promote interaction with 2-oxogluterate (T184A, N197A, K204A, T285A). Each mutant rescued the differentiation deficiency due to JMJD6 knockdown. These data strongly support the conclusion that JMJD6 enzymatic activity is not required for adipogenic differentiation.

A recent report demonstrates JMJD6 has kinase activity that is dependent on the JmjC domain and the poly-serine stretch (33). Although the authors did not probe the requirement for iron or 2-oxogluterate binding, the ability of JMJD6 to function in adipogenic differentiation in the presence of mutations that block iron or 2-oxogluterate binding or that disrupt the poly-serine stretch makes it unlikely that JMJD6 kinase activity is required.

### The JMJD6 AT hook-like domain is required for promoting adipogenesis in culture

Mutation of the JMJD6 AT hook-like domain prevented the ability of ectopically expressed JMJD6 to rescue the adipogenic differentiation deficiency caused by JMJD6 knockdown. The results suggest that this domain is important for differentiation, though the mechanism of its action remains unknown. The JMJD6 AT hook-like domain does not fit the consensus for a classical AT hook nor for an extended AT hook domain. Instead it seems to be a hybrid. While an AT hook is a DNA binding surface (53), the extended AT hook is believed to be an RNA binding domain (54). JMJD6 has been described as a functional player in both DNA and RNA transactions, including transcriptional activation, release of paused RNA polymerase II, and splicing, among other processes. Some reports suggest that JMJD6 binds to RNA (17, 54). Perhaps the hybrid AT hook structure is indicative of an ability to bind to both DNA and RNA.

The mutations we introduced in the JMJD6 AT hook-like domain affect the hinge between two α-helices and are not predicted to significantly change the protein structure (Fig. 6). Given the difficulties experienced expressing JMJD6 mutants, the subtle nature of the effect on structure by these alanine substitutions may explain why this mutant could be expressed and assayed. The main consequence of the mutation may be the elimination of the side chains of the mutated amino acids (GPRK; AA 303-307). Whether such mutations affect the ability of the AT hook-like domain to interact with its predicted nucleic acid substrates remains to be determined.

Multiple reports show JMJD6 interacting with genomic sequences via chromatin immunoprecipitation assays (10-14, 27), though this approach precludes determining whether or not JMJD6 binds to chromatin or binds indirectly through other factors that directly contact DNA. Interestingly, one factor implicated in JMJD6 interaction with chromatin is BRD4. Cooperativity between JMJD6 and BRD4 in transcriptional regulation has been shown (12, 69), and it appears that JMJD6 binding to chromatin is dependent on BRD4, while BRD4 binding is independent of JMJD6 (12). Whether the evidence for JMJD6 binding to chromatin indirectly applies to other regulatory roles or not remains to be determined. The connections between JMJD6 and BRD4, in adipogenesis as well as in other biological systems, suggest that the BRD4 interacting domain in JMJD6 would also be important for JMJD6 function. Our inability to stably express a JMJD6 protein with mutations in the BRD4 interacting domain may support that hypothesis if the interaction with BRD4 or the integrity of the BRD4 interaction domain is essential for JMJD6 functions and for long-term cell viability.

### Is JMJD6 a scaffold protein?

A requirement for JMJD6 in the absence of catalytic activity raises the possibility that JMJD6 also has scaffolding functions that facilitate molecular processes by serving as an interaction surface for many proteins. In fact, the number of JMJD6 interacting proteins is large, with binding partners implicated in transcription, splicing, and translation, among others (discussed above). The evidence that JMJD6 has myriad roles supports the idea that it may be more of a structural component of multi-protein complexes than a protein with a precise regulatory role. The ability to carry our demethylation and/or lysyl hydroxylation may contribute to the efficiency of some processes but may not be a strict requirement.

### JMJD6 domain requirements for oligomerization

One of the more curious observations about JMJD6 is its ability to oligomerize. The function of oligomerization is not understood. Some of the earlier observations determined that mutations in the JMJD6 iron binding amino acids abolished oligomerization (40). This led to the conclusion that catalytic activity was required for oligomerization. However, our results demonstrate that two of the single point mutations in JMJD6 amino acids that promote 2-oxogluterate binding did not oligomerize (K204A, T285A), while two mutants did (T184A, N197A). Since JMJD6 catalytic activity also requires 2-oxogluterate, the data invalidate the existing correlation between catalytic activity and oligomerization.

We also determined that there is no correlation between oligomerization and function in promoting adipogenic differentiation. All of the JMJD6 mutants that affected cofactor binding were functional in promoting JMJD6 function even though four of the six mutants were inhibited in the ability to oligomerize. Similarly, the mutations in the poly-serine stretch did not compromise JMJD6 function even though ability to oligomerize was compromised. In contrast, the mutations in the AT hook-like domain inhibited the ability of JMJD6 to promote adipogenesis even though the mutations had no impact on oligomerization. Based on the data, JMJD6 oligomer formation is neither required for nor correlated with adipogenic differentiation.

## Materials and methods

### Cell culture and in vitro differentiation

Mouse C3H10T1/2 (ATCC: CCL-226), HEK293T (ATCC: CRL-3216) and BOSC23 (ATCC: CRL-11270) were cultivated in Dulbecco’s modified Eagle’s medium (DMEM) high glucose medium (Life Technologies) containing 10% fetal bovine serum (FBS; Life Technologies) and 100 U/ml of penicillin/streptomycin (Life Technologies) at 37 °C and 5% CO_2_. Adipogenic differentiation of C3H10T1/2 cells was induced with DMEM containing 10% FBS, 0.5 mM 3-isobutyl-1-methylxanthine (IBMX), 1 µM dexamethasone, and 10 µg/ml insulin (all reagents from Sigma). After 2 days, the media was switched to DMEM, 10% FBS, 10 µg/ml insulin and changed every other day until harvested.

### Plasmid construction

Expression of either JMJD6 or scramble shRNA used lentiviral vectors as previously described (11, 70) Retroviral vectors for JMJD6 expression were generated by cloning the *Jmjd6* coding sequence with a C-terminal FLAG-tag sequence (11) into the pBABE plasmid vector containing the neomycin resistance gene (Addgene #1767). Mutant versions of JMJD6 were generated from the wild type construct using QuikChange Lightning site-directed mutagenesis kit (Agilent). Amino and carboxy terminal truncations of JMJD6 were generated by PCR amplification. The version of JMJD6 lacking the JmjC domain was generated by megaprimer-based PCR mutagenesis (71). All constructs were verified by DNA sequencing. Primers used for plasmid construction are presented in Table 1.

**Table 1.**
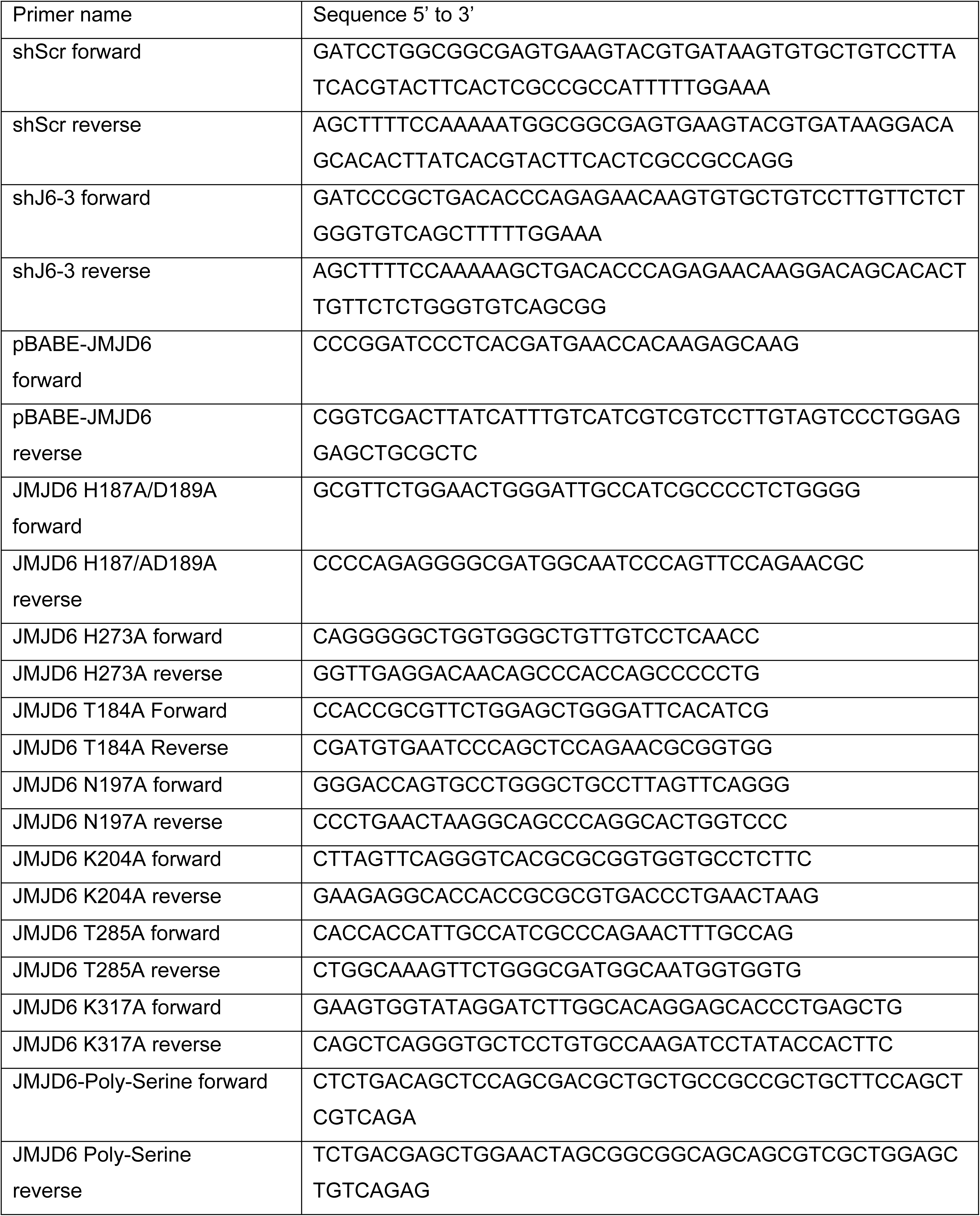
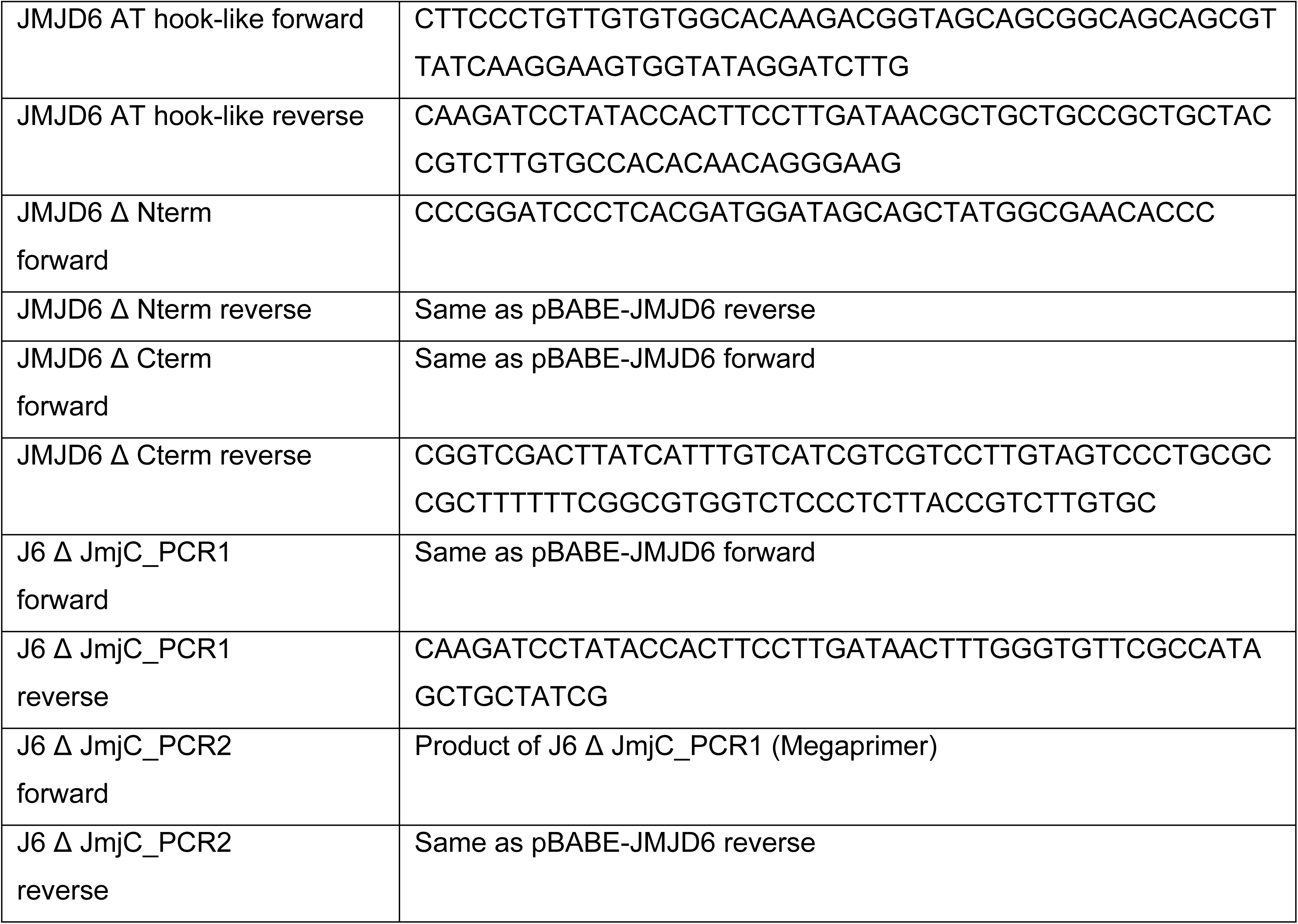
Oligonucleotides for plasmid construction.

### Virus production and transduction

Lentivirus production was performed as previously described (70). Briefly, the pLenti/sh*Jmjd6* or pLenti/shCtrl constructs were co-transfected with pLP1, pLP2 and pVSVG packaging vectors into HEK293T cells. Retroviral vectors expressing either wild type or mutant/truncated JMJD6 versions were individually transfected into BOSC23 cells (72). Transfection was performed using Lipofectamine 2000 according to the manufacturer’s instructions (Invitrogen). The medium was replaced the following day with DMEM containing 10% FBS and antibiotics; viral supernatants were harvested 48 h post-transfection and filtered through 0.45 µm syringe filter (Millipore). For viral transduction, 5 ml of the filtered supernatant supplemented with 8 µg/ml of polybrene (Sigma) were used to infect 3×10^6^ cells. After overnight incubation, transduced cells were selected with media containing 2.5 μg/ml puromycin (Invitrogen), or 500 µg/ml of G418 (Sigma) as needed.

### Oil Red O staining

Differentiated C3H10T1/2 cells were washed with PBS and fixed with 10% phosphate-buffered formalin (Fisher Scientific) overnight at room temperature. Subsequently, cells were washed with 60% isopropanol and air dried. Cells were stained with filtered 60% Oil Red O solution (AMRESCO) for 10 min and washed extensively with water. To quantify staining, Oil Red O was extracted using 100% isopropanol and the eluted dye was measured at optical density 500 nm.

### Western blot analysis

Cells were washed twice with ice cold PBS and harvested in RIPA buffer (50 mM Tris-HCl, pH7.4, 150 mM NaCl, 1mM EDTA, 1% NP-40 and 0.25% sodium deoxycholate) supplemented with Complete protease inhibitor cocktail (Roche). Protein concentration was measured using the Bradford protein assay following the manufacturer’s instructions (Bio-Rad). The samples were mixed with 6X loading buffer (240 mM Tris-HCl, pH 6.8, 8% sodium dodecyl sulphate (SDS), 40% glycerol, 0.01% bromophenol blue and 10% β-mercaptoethanol), and boiled at 95 °C for 5 min. Samples were separated on 10% SDS-PAGE and electro-transferred onto PVDF membranes (Bio-Rad). Then the membranes were blocked with 5% non-fat milk in PBS-0.1% tween for 30 min at room temperature, and the proteins of interest were detected by incubation with the specific primary antibodies diluted in 5% non-fat milk/PBS-0.1% tween overnight at 4 °C. Subsequently, the membranes were incubated with the species-appropriate HRP-conjugated secondary antibodies in 5% non-fat milk/PBS-0.1% tween for 2 h at room temperature. Chemiluminescent detection was performed with ECL PLUS (GE Healthcare). Primary antibodies used to detect JMJD6 (sc-28349; 1:200 dilution) and β-ACTIN (sc-81168; 1:2000 dilution) were purchased from Santa Cruz Biotechnology. FLAG-tagged proteins were detected using a 1:2000 dilution of a previously described polyclonal rabbit antisera against a peptide encoding the FLAG epitope (73). The HRP-conjugated secondary antibodies (NA9340 and NA9310) were purchased from GE Healthcare Life Sciences and used at 1:5000 dilution.

### Gene expression analysis

Total RNA was isolated using TRIzol reagent (Invitrogen), and treated with DNAse I (Invitrogen) according to the manufacturer’s instructions. cDNAs were prepared from 1 µg of total RNA with random hexamers (Roche) and the Superscript III reverse transcriptase kit (Invitrogen) following the manufacturer’s instructions. Changes in gene expression were analyzed by quantitative PCR using Fast SYBR-Green master mix (Thermofisher Scientific) using the QuantStudio 3 Real-Time PCR System (Thermofisher Scientific). Relative gene expression levels were calculated as 2(^Ct*Eef1a1* – Ct*gene*^) and were normalized to the experimental control as indicated (74). Primers for gene expression analysis are listed in Table 2.

**Table 2.**
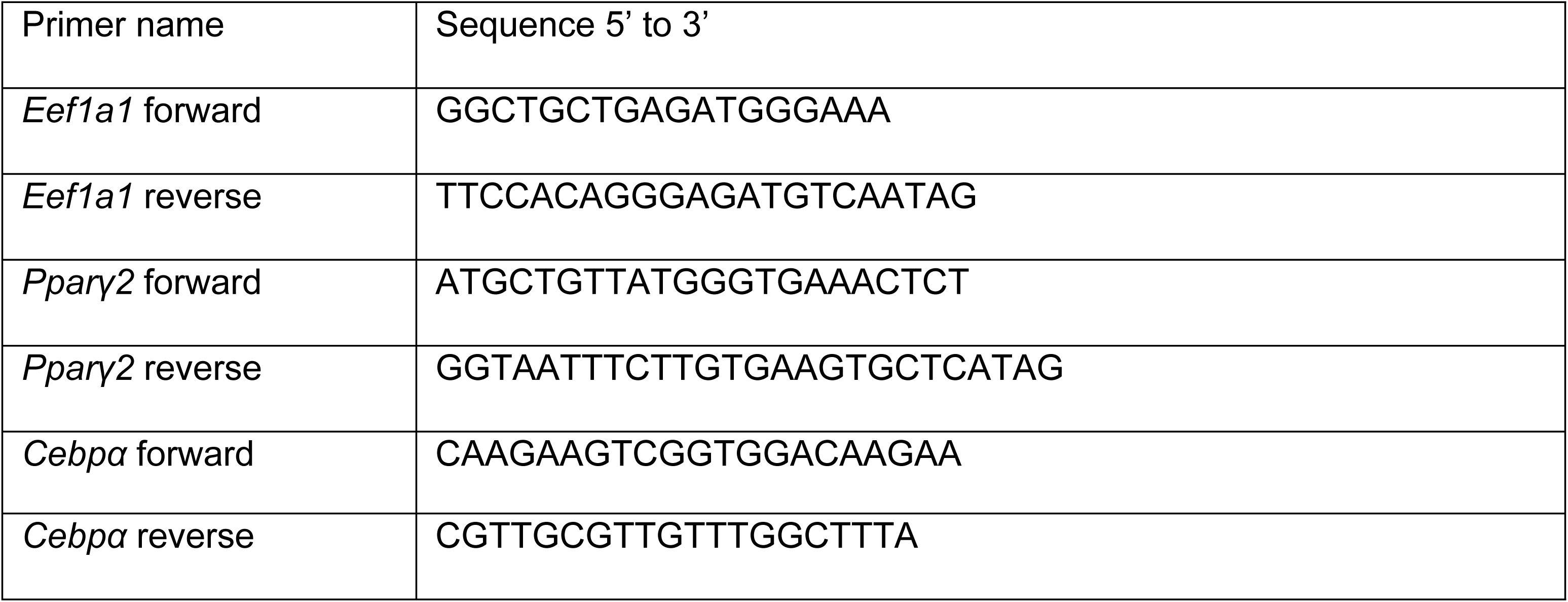
Primers for gene expression analysis.

### Statistical analysis

All quantitative data are shown as mean +/- the standard deviation (SD) of at least three independent biological replicates. Statistical analyses were performed using Student’s t-test with two-tailed distribution and equal variance. Significance is displayed with *P < 0.05 and **P < 0.01.

### Structure Modeling

The crystal structure of JMJD6 (ID 3K2O) was downloaded from The Protein Data Bank and generation of the image of the dimeric structure was made with the molecular visualization program PyMOL (Version 1.5.0.4; Schrödinger, LLC) using the entire PDB sequence. The structure of the portion of the AT hook-like domain that is depicted in Fig. 6 was generated by selection of amino acids 296 to 316. Subsequently, amino acids 303 to 307 were substituted for alanines to generate the structure of the mutant version.

## Acknowledgements

We thank Dr. S Syed for advice, Dr. H Witwicka for comments on the manuscript, and Dr. T Padilla-Benavides for advice and comments on the manuscript.

## Author contributions

PRG and ANI designed the project and experiments and analyzed data. PRG and JWCR performed experiments. PRG and ANI wrote the manuscript. ANI was responsible for funding acquisition.

